# Tactile Mechanisms and Afferents Underlying the Rat Pup Transport Response

**DOI:** 10.1101/2024.08.23.609194

**Authors:** Zheyi Ni, Connor Neifert, Arturo Rosete, Abdalla M. Albeely, Yu Yang, Marta Pratelli, Michael Brecht, Ann M. Clemens

## Abstract

Juvenile rodents and other altricial mammals react with calming, immobility and folding up of feet to parental pickup, a set of behaviors referred to as transport response. Here we investigate sensory mechanisms underlying the rat transport response. Grasping rat pups in anterior neck positions evokes strong immobility and folding up of feet, whereas more posterior grasping positions have lesser effects on immobility and foot position. Transport responses are enhanced by slow (1Hz) and even more so by fast (4Hz) gentle shaking and translation of the pup, features consistent with parental transport. In response to lateral grasping, the forepaw below the grasping position points downwards and the forepaw lateral to the grasping position points upwards and medially. Such forepaw adjustments put the pup’s center of gravity below the grasping point, optimizing pup transportability along with folding up of feet and tail lifting. Tactile stimuli on the back, belly, tail, whisker, dorsal forepaws and dorsal hind-paws do not significantly affect the behaviour of anterior-neck-held pups. Instead, ground contact or paw stimulation consistent with ground contact disrupts transport responses. We identify afferents mediating the transport response by examining membrane labelling with FM1-43 following anterior neck grasping. We observe a dense innervation of the anterior neck skin region (∼30 terminals/ mm^2^). We also observed an age-related decrease of cytochrome oxidase reactivity in the rat somatosensory cortical neck representation, a possible correlate to the developmental decrease in the pup transport response. We conclude anterior neck grasping and loss of ground contact trigger calming and postural adjustments for parental transport in rat pups, responses putatively driven from the densely innervated anterior neck skin.

## Introduction

Altricial mammals are born helpless and defenceless; thus, their physiology supports mechanisms for maternal care to support their survival. One such maternal protection mechanism is the transport response, where carrying behavior by the mother elicits a calming phenotype in infant humans and mice. In humans, walking transport helps babies to fall asleep (Esposito et al. 2013). In quadrupedal mammals, transporting offspring away from potential environmental threats or towards resources and safety is crucial though mothers must rely on grasping the body to move their infants (Esposito et al. 2019; Yoshida et al. 2013).

Earlier work showed that in mice transport response is associated with both pup postural adjustments (i.e. including compact posture, folding up of the feet and lifting of the tail) (Esposito et al. 2013), and calming effects (cessation of body movements and distress vocalizations, increased thresholds for pain responses and slowing of the heartbeat) (Esposito et al. 2013; Yoshida et al. 2013). While postural adjustments require the correct functioning of cerebellar circuits, pharmacological evidence points to parasympathetic contributions to calming effects (Esposito et al. 2013). The importance of these effects emerged clearly in experiments where pup transport response was abolished by local anesthesia, and as a consequence mothers took more time to retrieve their pups and bring them back to the nest (Esposito et al. 2019).

A seemingly simple behavioral stimulus like maternal pickup can evoke comprehensive postural, vocalization, autonomic, and nociceptive alterations in human infants and rodent pups. Despite being a highly evolutionary conserved and reliable behavioral response to tactile stimuli and movement, few studies have attempted to understand the mechanistic basis of the transport response.

## Results

To better understand the tactile mechanisms controlling the rat pup transport response we posed the following questions: (i) what skin regions drive the rat transport response? (ii) Does the precise mode of pup-carrying affect transport responses? (iii) Do pups react to how they are carried? (iv) Can the sensory afferents activated in the transport response be identified?

We analyzed the rat pup transport response in four steps. First, we investigated sensory stimuli triggering the transport response, then we investigated postural adjustments during transport as well as the stimuli that terminate the transport response. Finally, we identify mechanosensory afferents that putatively trigger transport responses.

### Anterior neck grasping induces the transport response

To investigate the tactile mechanisms mediating the response of rat pups to parental pickup, we tested which locus on the skin elicited a more pronounced transport response. We evaluated three distinct grasping positions around the pup’s neck: the anterior neck (a grasping position between the ears), the central neck (a grasping position between in the neck), and positions 5-10 mm posterior to the neck (Figure 1A). Since immobility is a hallmark of the transport response (Yoshida et al. 2013), we scored immobility according to the criteria described in Figure 1B. We found that grasping the anterior neck position is more effective in inducing immobility (Figure 1C, anterior vs central, p =0.0225; anterior vs posterior, p < 0.0001; central vs posterior, n.s.). Because pups also adjust posture in response to parental carrying by retracting their hind-paws and orienting them upwards (Yoshida et al. 2013), we scored foot position according to the criteria described in Figure 1D. Most pups fold up their feet when the anterior position is grasped, whereas the posterior position has a substantially smaller effect on the foot position (Figure 1E; anterior vs central, p = 0.0176; anterior vs posterior, p = 0.0764; central vs posterior, n.s.). The markedly different effects of anterior and posterior neck holding positions are also shown in Movie 1. Thus, grasping the anterior neck induces immobility and a folding up of the feet, i.e., the transport response.

**Figure 1.**
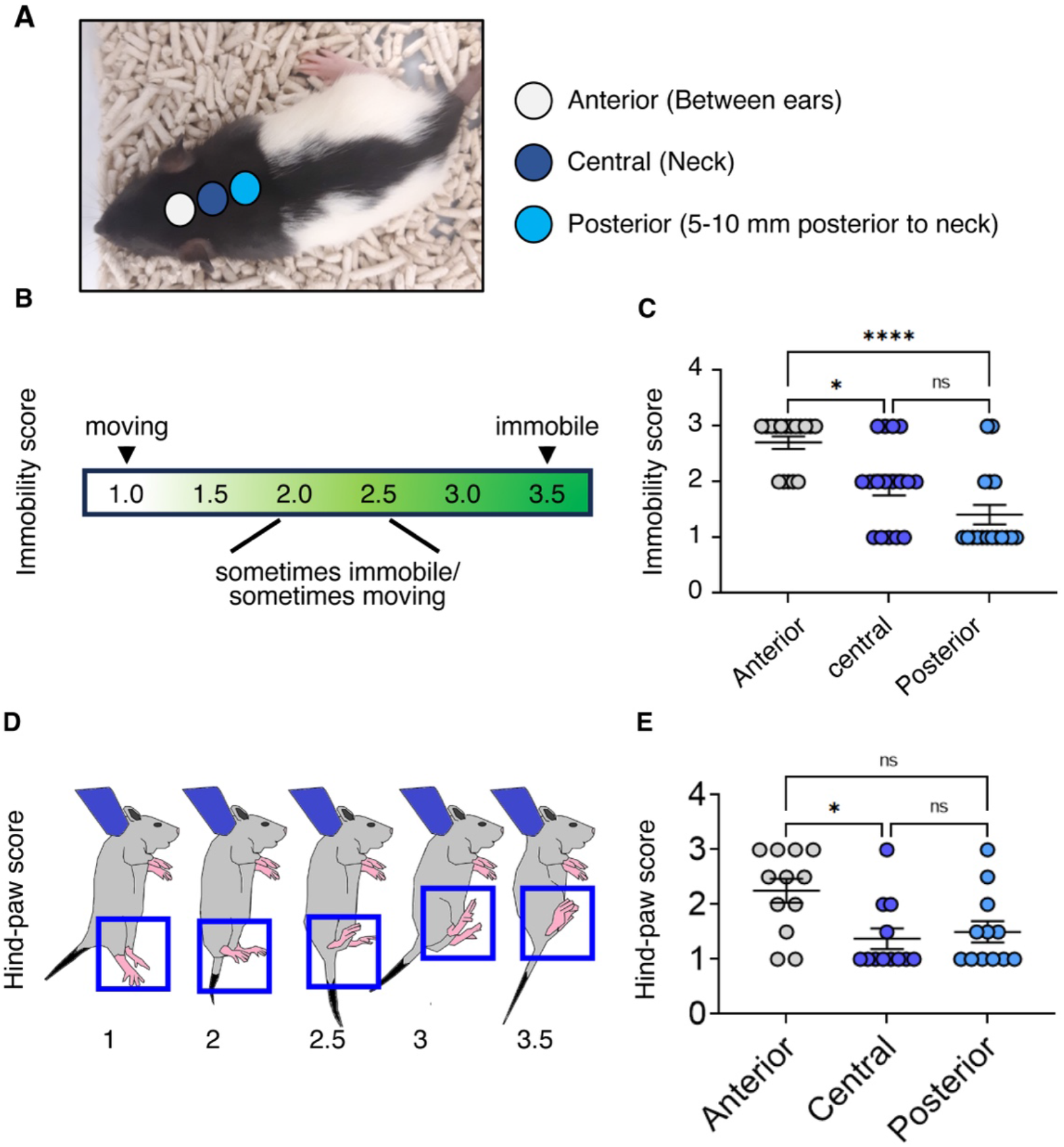
Grasping the anterior neck region evokes stronger transport responses than more posterior grasping positions. **A**. Rat pup with schematically superimposed neck grasping positions. **B**. The criteria for immobility score. **C**. The immobility score of grasping different parts of the neck. **D**. The criteria for hind-paw score. **E**. The hind-paw score grasping different parts of the neck. *p<0.05, ****p<0.0001. Statistical analysis was performed using Kruskal-Wallis test followed by Dunn’s multiple comparisons test. Associated Movie 1: Anterior vs Posterior Holding position. 25-day old rat pups are shown being held in the anterior position, with grip placed between the ears, for 12 seconds, the pups remain fully immobile. In contrast, when held in the posterior position, the rat pups demonstrated mobility (https://figshare.com/s/809990246af6a3c6ab45).

### Stimuli mimicking ambulation induce the transport response

Next, we wondered if mimicking parental ambulation would increase rat pup transport responses. Because the adult rat stepping rhythm has a frequency of 4Hz (Joshi et al. 2023), we included a 4Hz up and down gentle shaking among the conditions tested. We scored the transport response in pups that were either: held stationary, shaken gently at 1Hz, shaken gently 4Hz, as well as shaken gently at 4Hz while translated horizontally (Figure 2A). To determine the efficiency of each condition tested in inducing transport response, we assessed the immobility score (Figure 2B, Movie 2) and the hind-paw score (Figure 2C). Gentle shaking at 4Hz and horizontal translation induced the highest immobility scores compared to stationary condition (p = 0.007) and 1Hz shaking (p = 0.0497) (Figure 2B). No other differences were observed. Shaking at 4Hz showed a significantly higher hind-paw score compared to stationary positions (p = 0.0352) and shaking at 1Hz (p = 0.0209) (Figure 2C). Additionally, shaking at 4Hz and horizontal translation showed a higher hind-paw score compared to stationary positions (p = 0.0036) and shaking at 1Hz (p = 0.0019) (Figure 2C). These findings show that mimicking parental ambulation (4Hz shaking combined with horizontal translation) induce the strongest transport response in rat pups.

**Figure 2.**
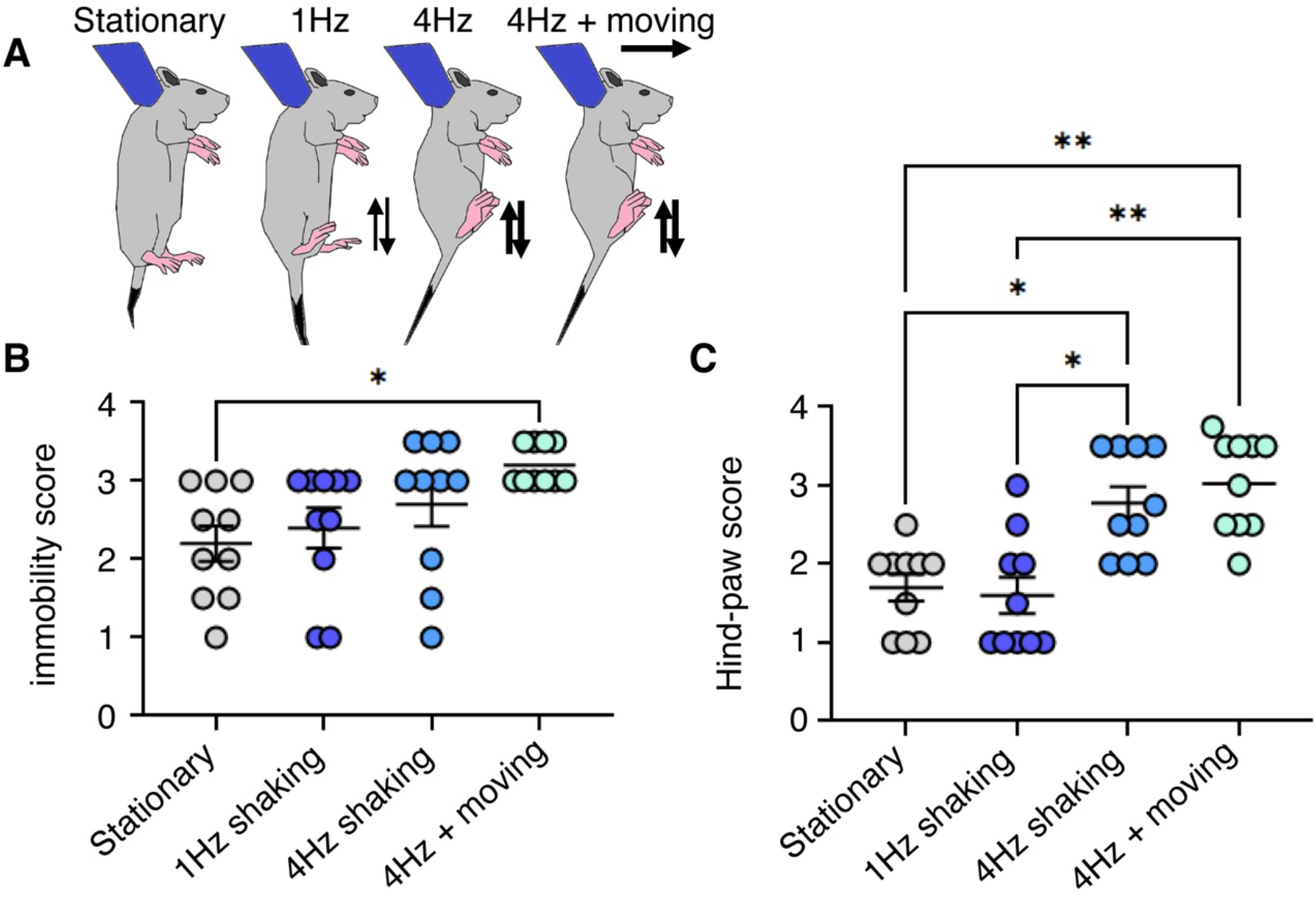
Gentle shaking and ambulation enhance transport responses in rat pups. **A**. Schematic of the experimental design showing P24 Long-Evans rats in the anterior hold position under stationary, shaking at 1Hz, shaking at 4Hz or shaking at 4Hz & moving conditions. Typical feet position during each condition also depicted. **B**. Immobility scores (see Figure 1) under stationary, shaking at 1Hz, shaking at 4Hz or shaking at 4Hz & moving conditions. No other significant effects were observed. **C**. Hind-paw scores (see Figure 1) under stationary, shaking at 1Hz, shaking at 4Hz or shaking at 4Hz & moving conditions. *p<0.05, **p<0.01. Statistical analysis was performed using Kruskal-Wallis test followed by Dunn’s multiple comparisons test. Associated Movie 2: 4Hz shaking induces the transport response. 25-day old rat pups were subjected to shaking at a frequency of 4 Hz for a duration of 12 seconds to simulate the maternal transport response. The pups were held in the anterior position and demonstrated sustained immobility throughout the stimulation period. (https://figshare.com/s/87bd2f87080aa4f28234).

### Laterally asymmetric grasping evokes systematic postural adjustments

Next, we assessed the transport response-induced postural adjustments when the grasping position was lateralized on either side of the neck. Holding either the left-anterior or the right-anterior side of the neck was surprisingly sufficient to induce limb contraction, tail lifting, and immobilization for each pup tested (Figure S1A). Interestingly, there was a differential, systematic postural change based on the side being held. In most animals, when the left-anterior side of the neck was grasped, the right forelimb was raised, and the left forelimb was lowered. The opposite was seen when the right side was grasped—the left forelimb was raised, and the right forelimb was lowered. There was a statistically significant association between the holding side and the arm raised (Figure S1B; p = 0.0286). This systematic difference in posture was seen across and within animals, suggesting that lateralized holds induce a conserved compensatory strategy.

### Tactile stimuli indicating ground contact terminate the transport response

We next sought to understand the factors terminating the transport response. To this end, we investigated the ability of various stimuli to disrupt the transport response. We focused on various tactile stimuli (ground contact, tail touch, belly touch, back touch, touch of one forepaw (1FP), touch of two forepaws (2FPs), touch of one hind paw (1HP), touch of two hind paws (2HPs)) (Figure 3A). We defined a transport reflex reversal score (the sum of decreases in immobility score (Figure 1B) and foot position score (Figure 1D). Ground contact drove a significantly higher reversal score compared to stationary hold (p < 0.0256) and tail stimulation (p < 0.0256) (Figure 3B). In separate experiments, we also tested whisker stimuli and observed little effects of whisker stimuli on transport responses. Transport reflex reversal scores also differed significantly between the 2FPs compared to stationary hold (p < 0.0022), tail (p < 0.0022), belly (p < 0.0441), back (p < 0.0105), and 1HP (p < 0.0105) stimulations (Figure 3B). The remarkable differences of one forepaw and two forepaws stimulation in potency to terminate the response between stimulating are also shown in Movie 3. Given that these initial results indicated that limb stimuli were critical to terminating the transport response, we chose to study the effects of paw stimulation more closely. We compared the effects of dorsal and ventral forepaw stimulation (Figure 3C). Ventral forepaw stimulation evoked a higher transport reflex reversal score than dorsal forepaw stimulation (p = 0.0012) (Figure 3D). Likewise, dorsal and ventral hind-paw stimulation was compared (Figure 3E). The ventral hind paw evoked a higher transport reflex reversal score compared to the dorsal hind-paw stimulation (p < 0.0006) (Figure 3F). Taken together, our data indicate that tactile stimulation that mimics the ground contact disengages the transport response in pups.

**Figure 3.**
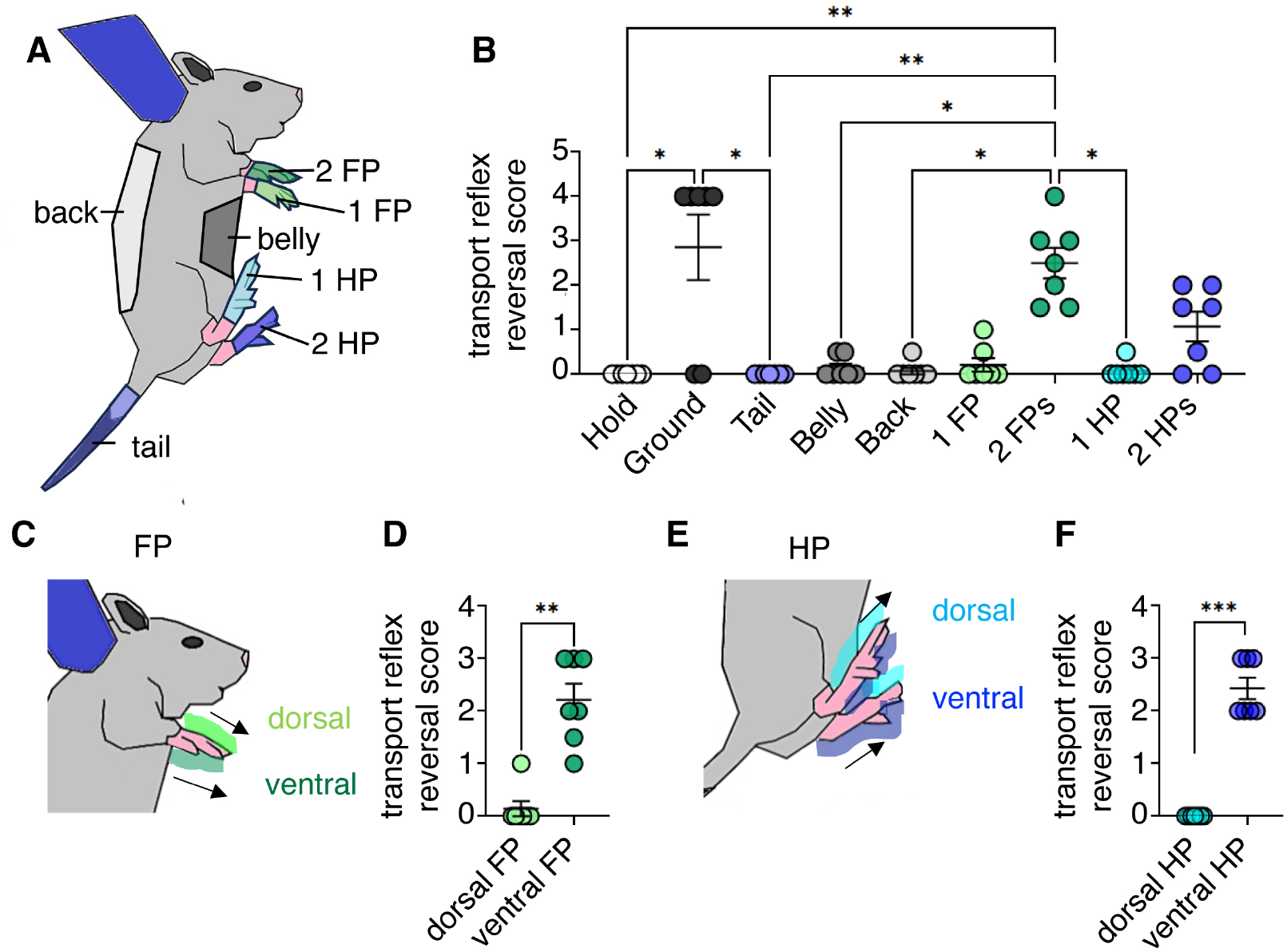
Tactile stimuli indicating ground contact terminate the transport response in rat pups. **A**. Schematic of P24 rat pup showing the anterior hold with the different stimuli location color coded. **B**. Transport response reversal score for different stimulation conditions (color codes as in **A**). The transport reversal score was as the sum of decreases of the immobility score and the hind paw score as defined in Figure 1. Ground contact, two fore-paw (2FP) stimulation and two hind-paw (2HP) stimulation disrupt the transport response. **C**. Schematic diagram of a rat pup showing the dorsal and ventral fore-paw stimulation. **D**. Ventral fore-paw stimulation evoked higher transport reflex reversal scores. **E**. Schematic diagram of a rat pup showing the dorsal and ventral hind-paw stimulation. **F**. Ventral hind-paw stimulation evoked higher transport reflex reversal scores. *p<0.05, **p<0.01, ***p<0.001. Statistical analysis was performed using Kruskal-Wallis test followed by Dunn’s multiple comparisons test (**B**) and two-tailed Mann-Whitney test (**D, F**). 1FP – one fore-paw; 2FPs – two fore-paws; 1HP-one hind-paw; 2HPs-two hind-paws. Associated Movie 3: Dual but not single fore-paw stimulation mobilizes the pup. 25-day old rat pup is shaken at 4Hz for 12 seconds. Stimulation of a single fore-paw does not disrupt the pup’s immobility. However, simultaneous stimulation of both fore-paws results in the pup becoming mobile (https://figshare.com/s/adb055f78e0670019789).

### Labelling of grasping-activated mechanosensory afferents in the anterior neck

We reasoned that activation of nerve terminals in the region where the mother grasps the rat is likely to trigger immobilization and changes in body posture during the transport response. Indeed, lidocaine-induced local anesthesia on the skin of the neck suppresses the transport response in mice (Esposito et al. 2019). To visualize mechanosensory afferents activated during the transport response we used the styryl dye FM 1-43. Due to its ability to penetrate cells via open ion channels and fluoresce when binding to intracellular membranes, FM 1-43 has been widely used to label sensory neurons (Meyers et al. 2003; Nishikawa 2011) and has recently been shown to be dependent on the activity of the mechanosensory PIEZO2 (Villarino et al. 2023). In the 24 hours that followed the injection with FM 1-43, rats were held by the anterior portion of the neck and subjected to five-minute-long sessions of shaking at 4 Hz (Figure 4A). The rat pups showed relaxation, immobility, and changes in body posture during each session for the entirety of the five-minute sessions. Samples of neck skin were collected 24 hours after FM 1-43 injection (Figure 4A). Dense FM 1-43 fluorescence was evident in the anterior but not the posterior portion of the neck (Figure 4B-F). In the anterior neck, we detected approximately 30 fluorescent dot-like structures per mm^2^ (Figure 4C, E), indicating the presence of sensory terminals associated with hair follicles. In contrast, reduced dot-like fluorescent structures were observed in the posterior region of the neck (Figure 4D, F), indicating a lack of activation of sensory afferents in this region. There was a higher density of fluorescent structures in the anterior portion of the neck than that in the posterior portion of the neck (Figure 4G; p = 0.0079). Overall, these data show that grasping activates mechanosensory terminals in the anterior portion of the neck, which are likely key afferents for initiating the transport response.

**Figure 4.**
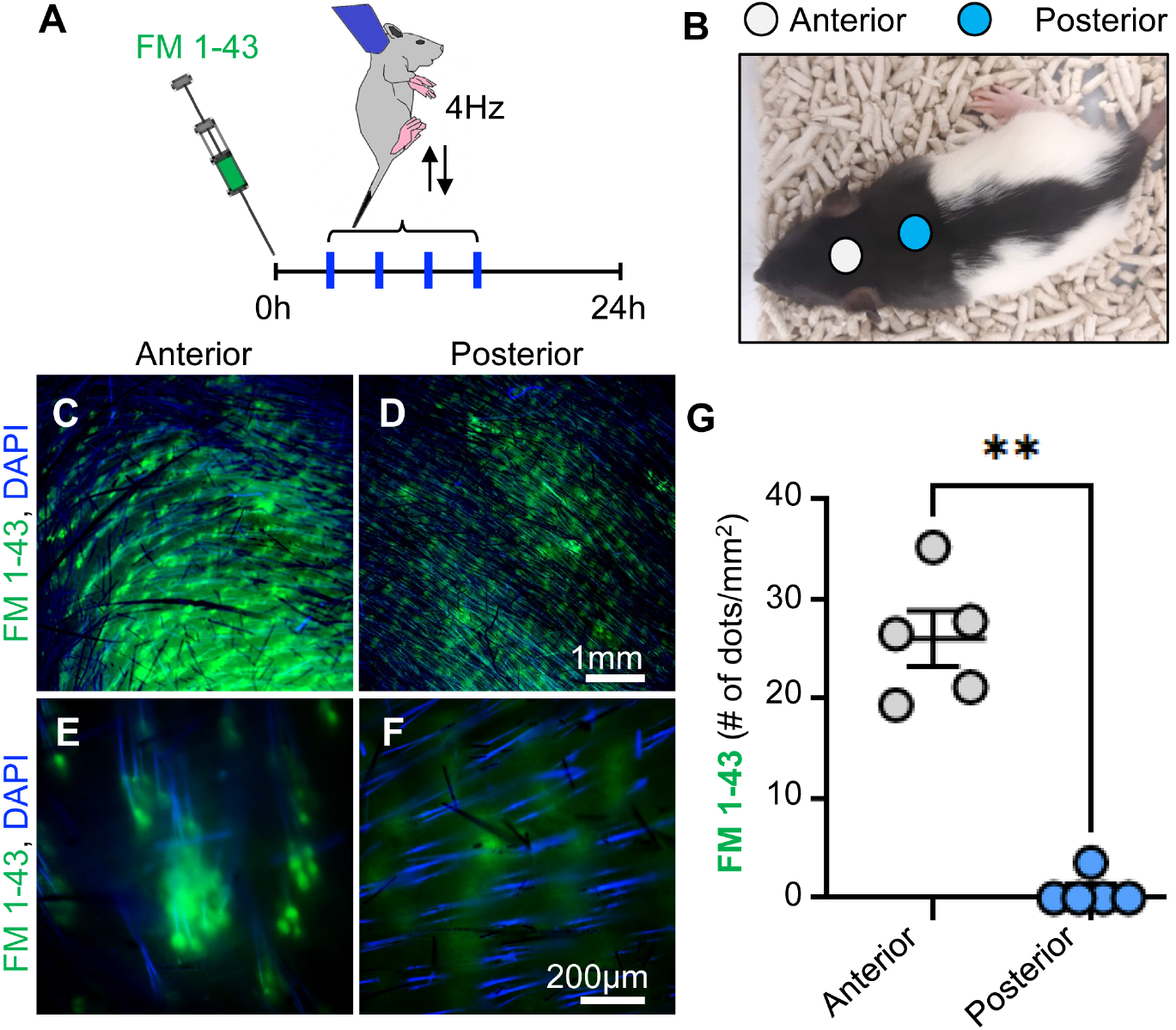
FM 1-43 labelling of anterior neck afferences activated by grasping. **A**. Experimental timeline. **B**. Representative image illustrating the neck “anterior” and “posterior” locations used for the analysis. **C, D**. Representative image FM 1-43 labelled follicles in the anterior (**C**) and posterior (**D**) portion of the neck. Scale bar 1mm. **E, F**. Representative high magnification image of FM 1-43 labelled dots in the anterior (**E**) and posterior (**F**) portion of the neck. Scale bar 200μm. **G**. Quantification of FM 1-43 labelled dots in the anterior and posterior portion of the neck. N=5 rats. **p<0.01. Statistical analysis was performed using a two-tailed Mann-Whitney test.

### Sensory responses and a loss of layer 4 thickness of somatosensory neck cortex in old rats

In the final part of our study, we investigated the neck region of the somatosensory cortex. To this end, we targeted coordinates (3 posterior, 3.5 lateral from bregma) for cellular recordings according to the body maps provided by Chapin and Lin (1984). Indeed, recordings at these coordinates returned a position consistent with the neck in the layer 4 somatosensory cortex body map (Figure S2A). We obtained whole-cell recordings at such coordinates in young rat pups and observed depolarization of the membrane potential following neck stimulation (Figure S2B). We also found that layer 4 of the somatosensory neck cortex is much thicker in young animals than in adult rats (Figure S2C). Indeed, the somatosensory neck cortex appears to be part of the somatosensory cortex with the most marked thinning of layer 4 with age (Figure S2D), as well as when juvenile layer 4 cortical thickness is normalized to adults one finds the thickest cortex in the somatosensory cortex neck region (Figure S2E). We conclude that layer 4 thickness in the neck region of the somatosensory cortex is greater in young juvenile animals and this declines with developmental age.

## Discussion

Here we show that grasping the anterior portion of the neck induced more pronounced immobility and postural adjustment which are boosted by mimicking parental ambulation via 4 Hz shaking and translation stimuli. Pups show systematic postural adjustments to lateral grasping. While pups in which the transport is activated ignore most stimuli, ground contact and ventral paw-touching stimuli systematically disrupt immobility and postural adjustments. FM1-43 labelling in combination with anterior neck grasping identifies rich afferent innervation of the anterior neck skin. We further show that the somatosensory representation of the neck area in juveniles is enriched and thicker than in aged animals. Our findings are in line with previous results which describe the physiological mechanisms of the transport response in mice (Esposito et al. 2013, 2019, Yoshida et al. 2013); we build upon this work by identifying the peripheral region responsible for the initiation of the transport response, the dynamics of the motion and temporal frequency required for its initiation.

### Anterior neck afferents and the transport response

Our findings indicate that the neck region of the rat is particularly sensitive to the initiation of physiological calming. Indeed, in humans massage techniques are typically aimed towards stimulation of the neck and head region as especially sensitive areas for calming effects (Personal communication Stephanie White, Sefton et al. 2011, Fazeli et al. 2016). We wonder how the calming effects of massage may be related to the transport response as we have described in juvenile rodents. Further examination of the density of mechanosensory afferents in the neck and head region across species may shed light on common mechanisms.

### Lateralized holds evoke systematic posture changes

The systematic postural changes seen during lateralized holds are likely to arise from the pups employing postural compensatory strategies to maintain balance. The rats in this study were aged P21-32, and it is understood that the rat vestibular system is fully developed and mostly matured by this age (Jamon, 2014). It is also well known that the righting reflex develops *in utero* to maintain upright posture (Sekulic, 2009). This systematic postural change may be attributable to maintaining balance: by extending the forelimb on one side, the pups may be offsetting the angle they are dangling from while being held from one side of the neck.

### Circuit understanding of the transport response

We performed in vivo whole cell patch clamp recording in the neck region of the somatosensory cortex of rat pups and identified age-dependent thinning of cytochrome oxidase staining layer 4 of the somatosensory cortex (Figure S2). These results in combination with the FM1-43 labelling of sensory afferents in the anterior region of the rat pup neck skin (Figure 4) indicate that specific circuits may mediate the tactile response which facilitates the rat pup transport response. Future work using circuit tracing methods from the skin sensory afferents to the somatosensory cortex should identify the route of the transport response circuitry. Further developmental studies should identify how the observed layer 4 thinning may occur. These insights will contribute more broadly towards our understanding of somatosensory development and circuitry which connects the body and the brain.

### Conclusion

A large body of research has focused on the maternal brain and how it is responsible for pup retrieval (Ehret 1992; Ehret & Koch 1989; Schiavo et al. 2020; Marlin et al. 2015; Dulac et al. 2014; Wu et al. 2014). Our focus is different, namely we examine how pup physiology is uniquely tuned for participation in maternal transport. Indeed, a mother transporting a calm offspring is an entirely different behavioral experience when compared with transport of an anxious and distressed pup. We identify region-specific mechanosensory afferents of the skin which contribute to a robust change of behavioral state which we relate to a natural behavioral phenomenon linking cooperative survival between parent and offspring.

## Methods

### Animals

Long–Evans rats (P21–P32) were used in these experiments. Rats were housed in a temperature and humidity-controlled environment. Experiments complied with regulations on animal welfare and were approved according to international law for animal welfare, ARRIVE guidelines and approved by the UK Home Office and the Woods Hole, USA Institutional Animal Care and Use Committee (23-09C and 24-09E).

### Transport response tests and videography

To test the transport response behavior of rat pups, experimenters gently grasped the neck region of rats while wearing powder-free nitrile examination gloves. Separation time from the litter was minimized and limited to the testing period. Filming was performed using a C920 Pro HD camera. Grasping areas, frequencies and transport configuration were signalled by event cards. The anterior grasping region was centered on the head region between the ears of the rat, central was defined by the neck region, and posterior was defined as 5-10mm posterior to the neck. The shaking frequencies were standarised using a metronome set to either 1 Hz or 4 Hz frequency. With translation stimuli, the speed of movement was standard walking speed (estimated at 1.4 meters per second).

### In vitro analysis of fur and skin patches

Rat pups were injected intraperitoneally with FM 1-43FX (Invitrogen: F35355) at a dose of 1.12 mg/kg of body weight in PBS 24 hours before sacrifice. Transport stimulation was given at 4Hz frequency in sessions of 5 minutes, 4 times in the 24 hours. 24 hours after FM 1-43 FX injection, rats were then transcardially perfused with phosphate buffer saline (PBS) followed by 4% paraformaldehyde (PFA) and the skin was removed and drop-fixed overnight while flattening. The skin was then imaged using a Zeiss Axio Zoom V16 and a Zeiss Axio Imager Z2 microscope. Regions for imaging were chosen based on coordinates for grasping targets of the skin in the transport response behavior. Exposure and acquisition parameters were standardised across images and analyzed in ImageJ/FIJI.

### In vivo patch clamp recording

Long-Evans rats were anaesthetized using urethane (1.4 g/kg i.p.). Animals were confirmed to be fully anaesthetized when there was no response to pinching of the paw or tail. After confirmation of full anesthesia, the head was secured with stereotaxic ear bars. Incised tissue was locally anaesthetized with lidocaine. A rectal probe monitored body temperature, and a homeothermic blanket (FHC, Bowdoinham, Me., USA) maintained it at 37 ± 0.5 °C. A craniotomy was made above the neck somatosensory cortex (3 mm posterior, 3.5 mm lateral from bregma). Electrodes entered the neck somatosensory cortex perpendicular to the cortical surface. To establish the neck receptive field, puffs of air were presented to the neck region. Air puffs were generated from pulses of compressed air, delivered by a computer-triggered airflow controller (Sigmann Electronics, Germany).

### Cytochrome oxidase staining

Animals were deeply anaesthetized with an additional dose of urethane and perfused transcardially with prefix, followed by 4% PFA. Brains were removed, hemispheres were separated, and cortices were flattened between two glass slides separated by clay spacers. Glass slides were weighed down with small ceramic weights for ca. 3 hr. Afterwards, flattened cortices were stored overnight in 2% PFA and 100 μm tangential sections were cut on a vibratome. Sections were stained for cytochrome-oxidase reactivity using the protocol of Wong-Riley (1979).

### Statistics

Statistical analysis was performed using GraphPad/Prism or MATLAB (MathWorks, Natick, Massachusetts, USA). Non-parametric tests were performed if normality criteria were not satisfied, otherwise parametric testing was performed. Data were expressed as the mean ± the standard error of the mean (SEM) unless otherwise noted.

## Acknowledgements

This work was supported by the Marine Biological Laboratory (MBL), the Neural Systems & Behavior Course (NS&B), a training grant from the NIMH (R25MH059472), the Simons Initiative for the Developing Brain (SIDB), Humboldt Universität zu Berlin, the Bernstein Center for Computational Neuroscience Berlin, the German Federal Ministry of Education and Research. Ann Clemens is supported by the University of Edinburgh and a Simons Edinburgh Scientific Academic Track (Simons ESAT) fellowship. We thank Alberto Pereda, Stephanie White, Rosalie Maltby and the NS&B forever people.

## Supplementary information

Movie 1: Anterior vs Posterior Holding position. https://figshare.com/s/809990246af6a3c6ab45 25-day old rat pups are shown being held in the anterior position, with grip placed between the ears, for 12 seconds, the pups remain fully immobile. In contrast, when held in the posterior position, the rat pups demonstrated mobility.

Movie 2: 4Hz shaking induces the transport response https://figshare.com/s/87bd2f87080aa4f28234 25-day old rat pups were subjected to shaking at a frequency of 4 Hz for a duration of 12 seconds to simulate the maternal transport response. The pups were held in the anterior position and demonstrated sustained immobility throughout the stimulation period.

Movie 3: Dual but not single forepaw stimulation mobilizes the pup https://figshare.com/s/adb055f78e0670019789 25-day old rat pup is shaken at 4Hz for 12 seconds. Stimulation of a single forepaw does not disrupt the pup’s immobility. However, simultaneous stimulation of both forepaws results in the pup becoming mobile.

**Figure S1.**
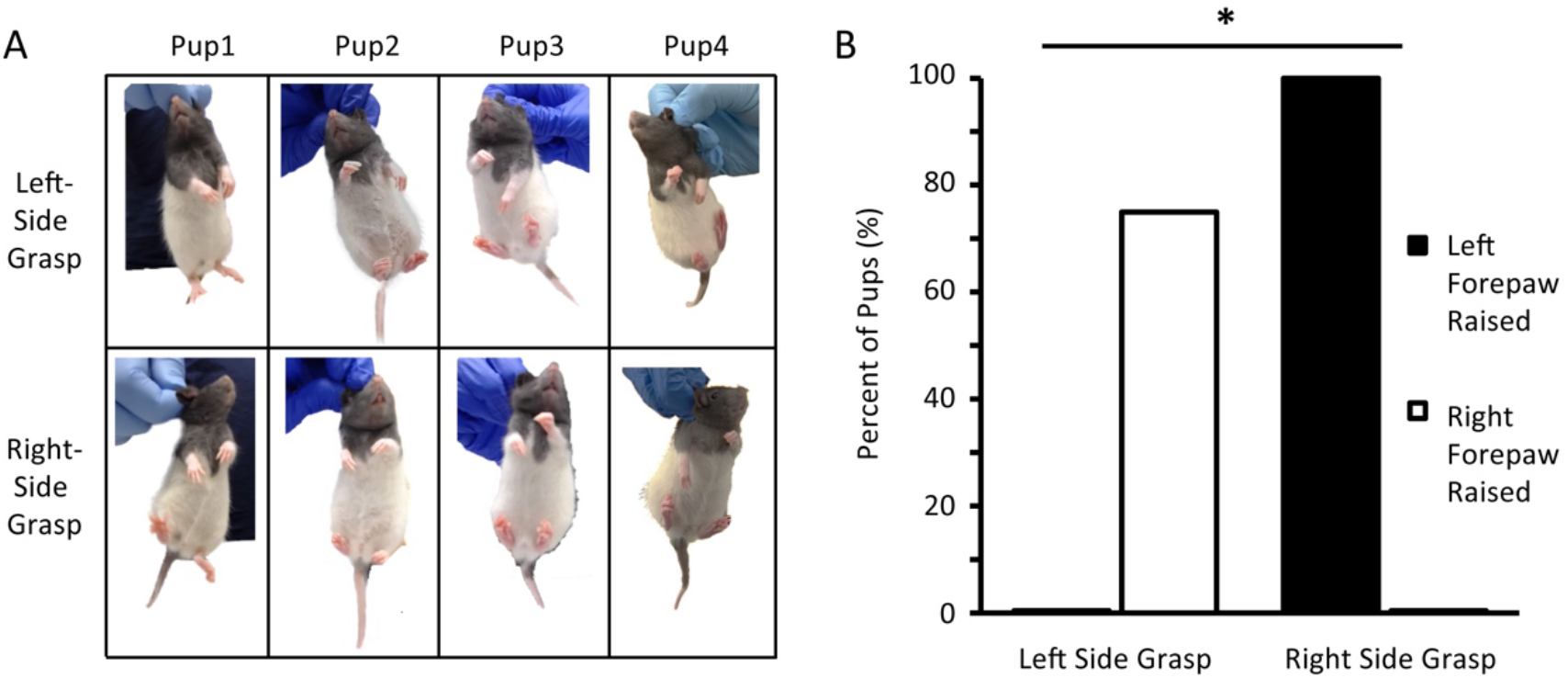
Laterally asymmetric grasping evokes systematic postural adjustments. **A**. Upper: holding only the left-side of the scruff results in raising of the right forelimb and lowering of the left forelimb. Lower, the reverse is seen when the right-side scruff of the same animal is held (N=4 rats). **B**. Asymmetric arm-raising behavior of pups. Percent is calculated as the number of pups exhibiting the behavior of to the total number of animals. * p < 0.05. Statistical analysis was performed using Fisher exact test.

**Figure S2.**
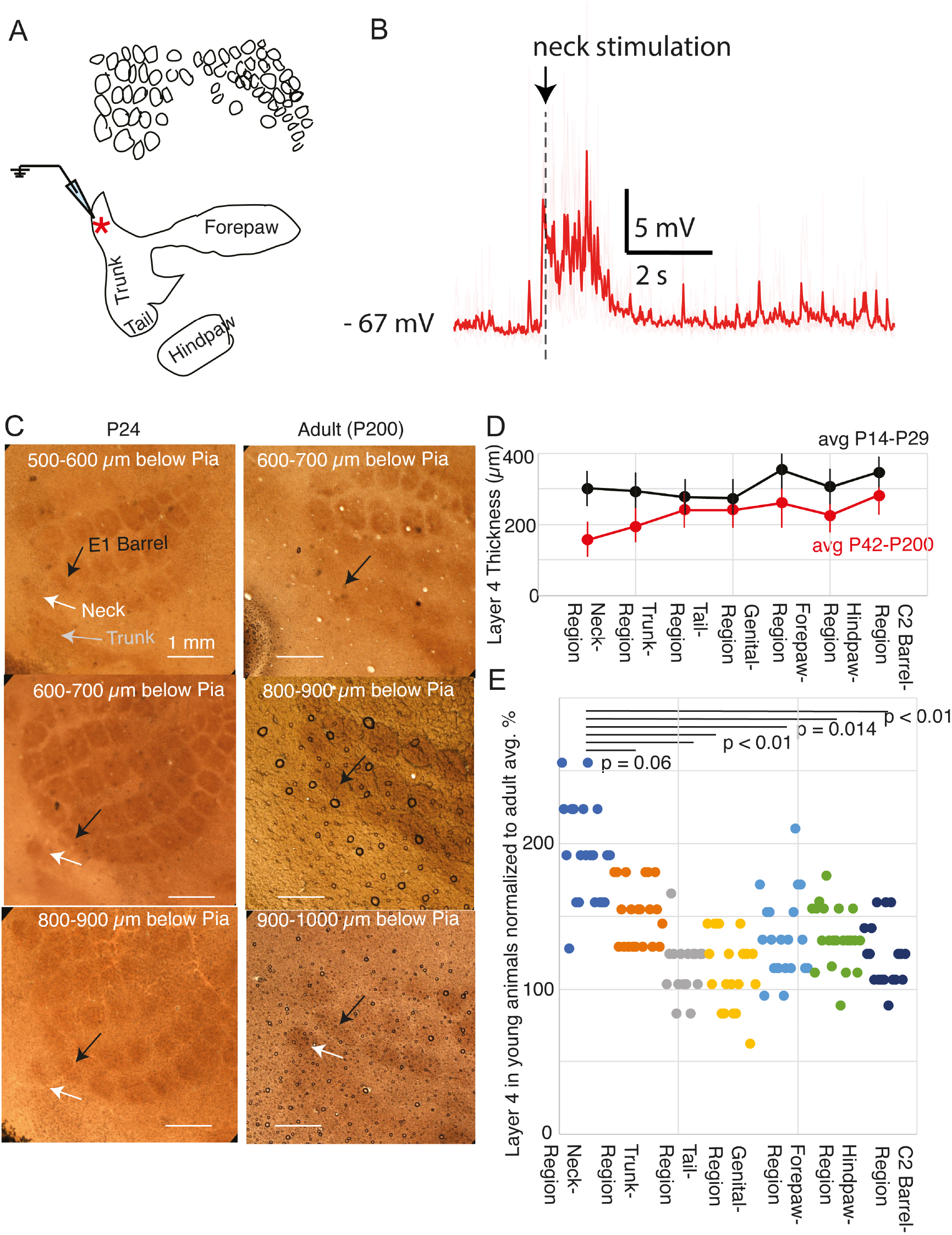
Responses of in rat somatosensory neck cortex and a thinning of somatosensory neck cortex from young to old rats. **A**. Line drawing of layer 4 body part contours from a cytochrome oxidase reactivity-stained tangential section through somatosensory cortex of a P24 rat pup, in which we obtained a whole-cell recording of a neuron at the site of the red star; the site was marked by an electrolytic lesion. **B**. Averaged (x5) voltage response to air puffs applied to the neck region from one representative cell in neck somatosensory cortex for a P23 rat pup. The stimulus is a sequence of air puffs at a rate of 4 Hz and comes on at second 2 and continues for 15 s (longer than shown). The responses are clear and habituate. **C**. Loss of cytochrome oxidase reactivity in neck somatosensory from young to old animals. Neck somatosensory cortex layer 4 (white arrow, identified as a dark contour in the cytochrome oxidase reactivity-stained tangential section through somatosensory cortex) can be identified through three 100 μm sections in young animals (left column). The same holds for the E1-barrel (black arrow) in young animals (left column). In adult animals (right column), however, neck somatosensory cortex layer 4 (white arrow, bottom) can be identified only in one 100 μm section. This differs from the E1-barrel (black arrow) in adult animals (right column), which can be identified in three sections. Scale bars are 1 mm. **D**. Average layer 4 thickness (identified by cytochrome oxidase reactivity) in different regions of somatosensory cortex; error bars refer to standard deviation (SD) in pups (black) and adults (red). **E**. Layer 4 thickness (identified by cytochrome oxidase reactivity) in individual somatosensory cortex maps normalized by average adult layer 4 thickness in the respective region. Neck somatosensory cortex stands out in relative thickness to adults.

